# Phylogenies can be used to predict future diversification but with limited success

**DOI:** 10.1101/2025.10.06.680809

**Authors:** Grace I Ridder, Jan Smyčka, David Storch, Arne Ø Mooers, Sarah P Otto

## Abstract

Understanding the processes leading to the emergence of new lineages is one of the major tasks of macroevolution as a scientific field. Multiple commonly used models and metrics allow to estimate the rates of species diversification at the tips of phylogenetic trees, and are thus potentially useful for predicting future evolutionary success of individual extant lineages. However, the predictive performance of these methods has never been tested with real-world data. Here we use a new approach to quantify the relationship between predictor tip rates and subsequent diversification rates derived from a probability distribution of numbers of daughter species. We explore if and how well the tip-rate methods predict near-future diversification using temporal slices across large fossil and extant tetrapod phylogenies. We find that tip rate estimates from these methods do predict diversification in all four extant and one out of three fossil phylogenies. The quality of these predictions varies across the tested methods, with the model-free approach being more robust overall but likelihood-based models delivering correctly scaled diversification rates. Our results suggest that future cladogenesis can be predicted using existing tip rate methods, but only moderately well and likely dependant on factors that are difficult to evaluate in practical applications.

## Introduction

Species diversification, the process of formation of biological diversity through speciation and extinction, runs at different paces in different evolutionary lineages [1–3]. Understanding this variation is crucial for inferring processes of the evolutionary past, explaining current patterns of species diversity, and predicting the fate of biodiversity in the future. Over the last several decades, researchers have developed an impressive toolbox of methods for capturing and exploring the variability of species diversification across time-calibrated phylogenetic trees [4–6]. These advances have been pivotal for associating different diversification rates with particular lineage characteristics, e.g. to identify links between species diversification and the appearance of key evolutionary innovations [6], various traits [7] or geographic features [8–10]. Similarly, there have been efforts, both qualitative and quantitative, to extrapolate our estimates of species diversification into the future [11,12] and use them for evaluating lineages for conservation planning [13,14].

Concomitant with wide usage of these methods is an ongoing debate about the appropriateness and mutual compatibility of the assumptions they make about the diversification process [5,15,16]. For example, the three methods commonly used for estimating rates of species diversification at the tips of phylogenies (so called tip rates) make fairly different assumptions about the variation of diversification among the lineages (Fig. 1). For instance, BAMM [6] assumes that diversification rates occasionally and abruptly change through evolutionary time, with the rest of the variation in evolutionary rates being less pronounced and changing only gradually. An alternative approach called ClaDS [5] assumes that the diversification rates change at every cladogenetic event but are still auto-correlated across the adjacent edges of the phylogeny. The commonly used model-free DR metric [4] makes no explicit assumptions about how variation arises, and simply notes that species diversification rate can be approximated by the inverse of a particular species isolation index [17], such that lineages with longer edges leading to the root have lower realized diversification.

**Figure 1:**
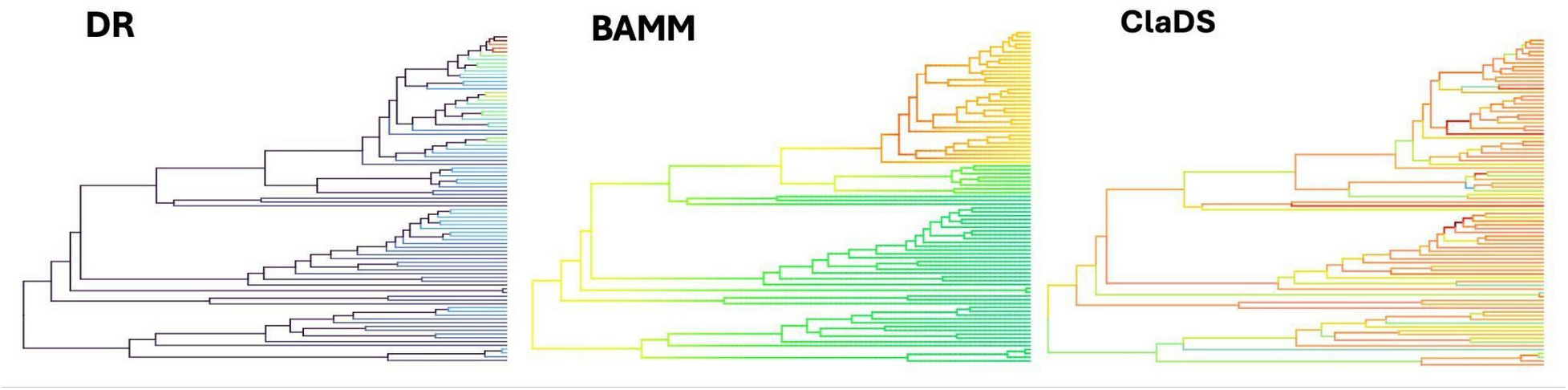
Illustration of how different tip rate methods interpret speciation rate variability on a single simulated phylogenetic tree. DR, which approximates speciation rates as inverted values of species isolation, progressively downweighted by distance from each focal tip, was shown to converge to the maximum likelihood estimates of speciation rate on pure birth trees [4]. BAMM assumes that speciation rates rarely punctually change during evolutionary time, and the rest of the variation in the rates is gradual [6]. ClaDS assumes that the speciation rates change at every cladogenetic event but are still auto-correlated across the tree [5]. Warmer colours denote faster estimated speciation rate.

A problem arises when these methods do not give compatible results. In such cases, the lack of external validation makes it impossible to untangle which of the assumed processes driving diversification variation are more appropriate, and so it is impossible to determine which results are the more credible. Even if the methods give similar estimates, which is fortunately often the case in empirical scenarios [8,15,18], accuracy and statistical performance of these estimates is still unknown. Previous simulation studies [5,15] explored errors associated with BAMM, ClaDS and DR on phylogenies generated using various forms of rate variability, such as gradually evolving rates, multiple rate regime shifts, and diversity dependence. They found that all the tree methods have non-zero predictive capacity in most scenarios. However, it is uncertain how well the tests on simulated phylogenies correspond to performance on empirical ones. The mechanisms they implemented in the simulations use ideas about rate variability common in the field, but are essentially the same ones that the tested diversification methods assume. A test using the full complexity of empirical data is thus needed to evaluate the real-world performance of these methods and adequacy of the evolutionary mechanisms they assume.

A potentially powerful empirical test of these diversification methods would explore whether their tip diversification estimates at one point in time accurately predict the diversity that individual tips generate subsequently. This is, of course, not possible in the present, as we do not have any information about the future. However, an approach presented by Cantalapiedra et al. [13] used phylogenies containing fossilized extinct species, calculated tip rates at arbitrary time slices and considered the “future” time as an arbitrary subsequent time bin. In their conservation-oriented study, they found that prioritizing species with the highest DR metric in the past led to slightly higher total diversity in subsequent time bins. This finding suggests that the DR metric has some predictive capacity, but the paper did not attempt to quantify this nor did it compare the predictive capacity of DR against other tip rate estimation methods. Another important limitation of their study was the exclusive reliance on fossil phylogenies. While they represent a data type that can be comfortably analyzed in a through-time manner, the fossil phylogenies are an incomplete representation of the past diversification process, and are known to be biased by nonrandom geographic distribution of fossil beds [19], different fossilization probabilities among taxa [20], and other related effects.

In this paper, we perform cross-validation tests of commonly used tip diversification methods (BAMM, ClaDS and DR) on temporal slices of both fossil and extant phylogenies (Fig 2). To do this, we developed an approach that mimics a generalized linear model, allowing us to link the tip estimates of diversification rates at a time slice with distributions of the number of subsequently surviving daughter species. This approach allows us to evaluate the relationship between estimated tip rates and realized rates of diversification in the subsequent slices of the tree, giving statistically meaningful and comparable measures of predictive capacity, such as generalized R^2^. Importantly, because we can condition the probability distribution of daugther species on survival of at least one lineage, our approach can be used with both fossil and extant phylogenies. Both these data types have important advantages and constraints - the fossil phylogenies are typically smaller, provide an incomplete and nonrandom representation of the past diversification, but do carry explicit information about extinctions; the extant phylogenies are more representative, but contain only strongly censored information about extinctions. Analyzing fossil and extant phylogenies in parallel should allow us to exploit the advantages of each data type and draw conclusions that are less dependent on their limitations.

**Figure 2:**
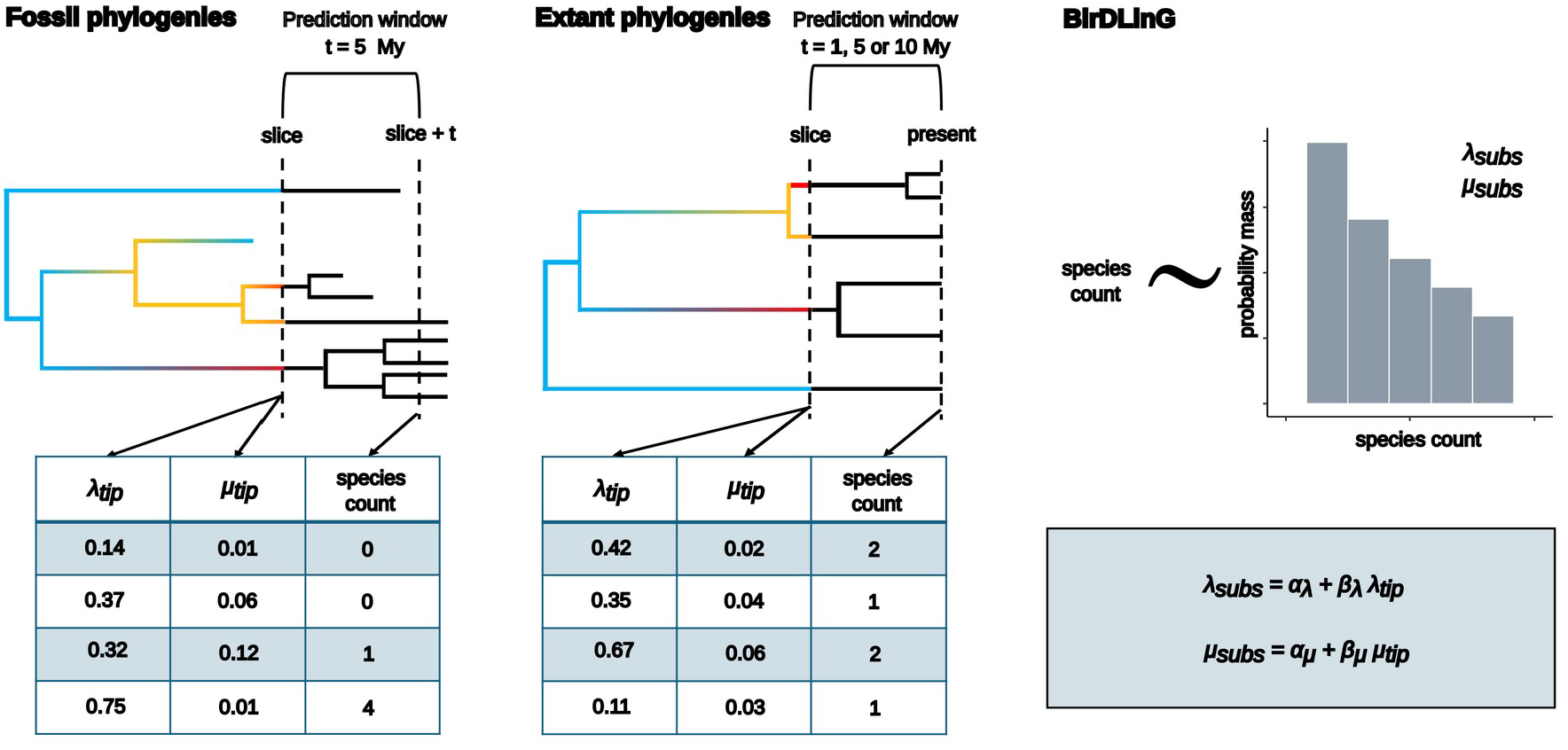
Schematic representation of temporal cross validation methodology used in this study for fossil (left) and extant (center) phylogenetic trees. The phylogenetic trees were sliced at a certain point of time, representing “present” in the cross validation test. The part of phylogeny preceding the slice, representing “past”, was used to estimate tip rates of speciation and extinction for all branches at the slice using BAMM, ClaDS or DR. For each branch identified at the slice, the number of subsequent daughter species after the prediction window was counted, representing the “near future”. Note that this daughter species count can be 0 for fossil phylogenies, but is >0 for extant phylogenies. Subsequently the daughter species counts were linked to the tip estimates of diversification using a likelihood based model (right, BirDLinG). This model assumed that numbers of daughter species were drawn from a distribution reflecting a homogeneous birth-death process with speciation and extinction parameters controlled by linear transforms of tip estimates of speciation and extinction rates for each tip at the slice.

## Materials & Methods

### The cross validation test linking tip rate estimates with subsequent diversity

In our cross validation test, we used extant and fossil phylogenies and sliced them at particular points in time, dropping the lineages after this point (Fig. 2). Each slice time thus represents the “present” time in which we perform the prediction in our test. On these truncated trees, representing the “past”, we employed the tip rate methods (BAMM, ClaDS and DR) to estimate diversification rates for each tip lineage at that slice time. For BAMM and ClaDS we estimated both speciation and extinction rates, and we considered DR to be an approximation of speciation rate [4] and its extinction rate estimates to be 0. To represent the “near future”, we counted the number of surviving daughter species from each tip after a subsequent prediction window of length *t*. The length of the prediction window was 5 My for fossil phylogenies, and 1, 5 and 10 My for extant phylogenies, where slices were positioned so that the end of prediction window was always at 0 My BP. Then we linked the tip rate estimates at the slice point and the subsequent number of daughter species using a custom developed likelihood-based approach called BirDLinG (standing for Birth-Death Linear Generalisation, [link to GitHub in accepted version]).

BirDLinG assumes that the number of daughter species descending from a tip in a subsequent time bin of length *t* can be locally approximated by a time-homogeneous birth-death process with speciation rate *λ* and extinction rate *μ*. The probability distribution of the number of surviving tips *n* on a fossil phylogeny resulting from a time-homogeneous birth death process after a prediction window of length *t* is

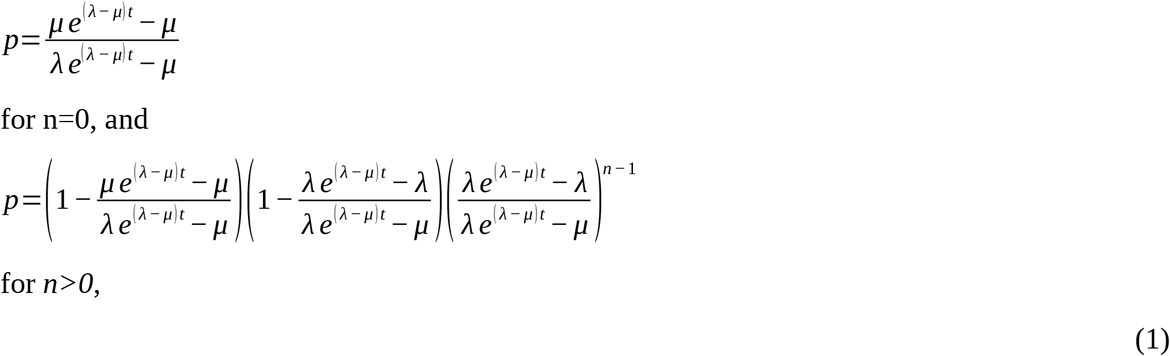

as described in [21]. Treated as a function of subsequent rates *λ* and *μ*, the Equation 1 serves as the likelihood of these parameters given the observed number of daughter species descending from a tip after time *t*. The total likelihood across all tips of one phylogeny slice can be consequently expressed as a product of such individual tip likelihoods. In BirDLinG, we assume that the vectors of subsequent speciation and extinction rates for all tips of one slice *λ_subs_* and *μ_subs_* are linear transformations of speciation and extinction rates estimated by tip rate methods *λ_tip_* and *μ_tip_*, so that

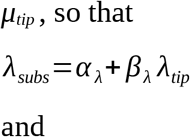

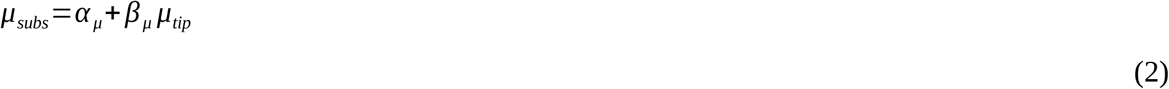

where the values of *α_λ_*, *α_μ_*, *β_λ_*, *β_μ_* are obtained using numerical likelihood optimization. With this formulation, we thus assume that there is a linear relationship between the tip estimates of diversification at the slice and the actual diversification rates generating the numbers of daughter species subsequent to the slice, and the direction of this relationship is controlled by slope parameters (*β*). This formulation accounts for the possibility that some of the tip rate methods systematically misestimate the subsequent diversification rates by a constant (*α*) or a proportion (*β*) but remain a linear transform of these values, which is still sufficient for many macroevolutionary applications. Also, formulated this way, our model mimics the structure of a generalized linear model (GLM), where the response variable *n* is drawn from the distribution defined by Equation 1, while the optimal parameters of this distribution *λ_subs_*and *μ_subs_* are latent and obtained via linear regression from a set of predictors *λ_tip_* and *μ_tip_*.

For the slices of extant phylogenies, our model formulation followed the same logic as above, but we conditioned the probability distribution in Equation 1 to contain at least one species (*n>0*), accounting for the fact that branches with no surviving offspring lineages would not have been observed as a tip in the sliced tree

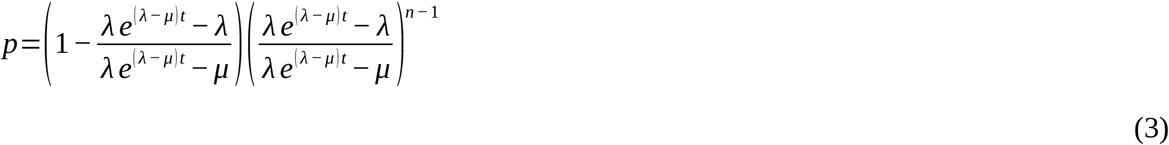

Importantly, this conditioning only works properly if the end of our prediction window (*slice+t*) is at 0 My BP from which the extant molecular trees were constructed (Fig. 2). Putting the end of the prediction window earlier in time while using the extant phylogenies would create a problem with lineages that survived to the end of the prediction window but went extinct later - such lineages are censored from the extant phylogenies which could not be properly accounted for. This limitation prevented us from slicing the extant phylogenies in a similar through-time manner as the fossil trees, so we can focus here only on slices in the last 1, 5 and 10 My BP, with all prediction windows ending at 0 BP. Another limitation of using the extant phylogenies lies in the fact that censoring 0 values from species counts effectively removes the majority of information about extinctions from our response variable. We still use the extinction component of BirDLing for exploring the extant phylogenies rather than forcing *β_μ_* to 0. We do this because the predictor tip extinction rates might still be, in theory, meaningful, and future species counts depend on net diversification rates rather than only speciation rates. However, we are aware that the estimates of these parameters are not interpretable on their own, and we also provide results for model with *β_μ_=0* in the supplementary materials (Supplementary Figure 1).

It is important to point out that in BirDLinG, we ignore the further heterogeneity in the birth-death process among the daughter lineages diversifying from any one tip in the prediction window subsequent to the slice, which is a relatively strong and potentially problematic assumption. Although this assumption follows the intuitive interpretation of tip rates in conservation biology [4,13], it, technically speaking, is at odds with the assumed rate heterogeneity before the slice. For the parametric methods (ClaDS and BAMM), it would be theoretically possible to implement the prediction window cladogenesis to follow the pre-slice processes, using frameworks like Generative Bayesian Models [22]. However, this approach would prevent us from testing DR, which, unlike ClaDS or BAMM, does not mechanistically deal with the pre-slice cladogenesis. We address this issue by two strategies: First, we only slice the phylogenies so that the pre-slice part of the tree is at least 3 times longer than the prediction window, allowing us to conclude that the prediction window is short enough to be approximated by a constant rate process. Second, we replicate our BirDLinG results with a Spearman rank correlation coefficient. This approach delivers very similar results (Supplementary Figure 2 and 3), and the squared Spearman’s Rho can be interpreted in a similar way as the generalized R^2^ values developed below. The Spearman correlation omits specific assumptions of BirDLinG, such as time homogeneity of the birth-death process in the prediction window and the linear relationship between tip estimates and true diversification rates. However, this generalisation comes at the expense of a parametric understanding of the system and also the potential to integrate tip estimates of speciation and extinction into a single predictive measure.

**Figure 3:**
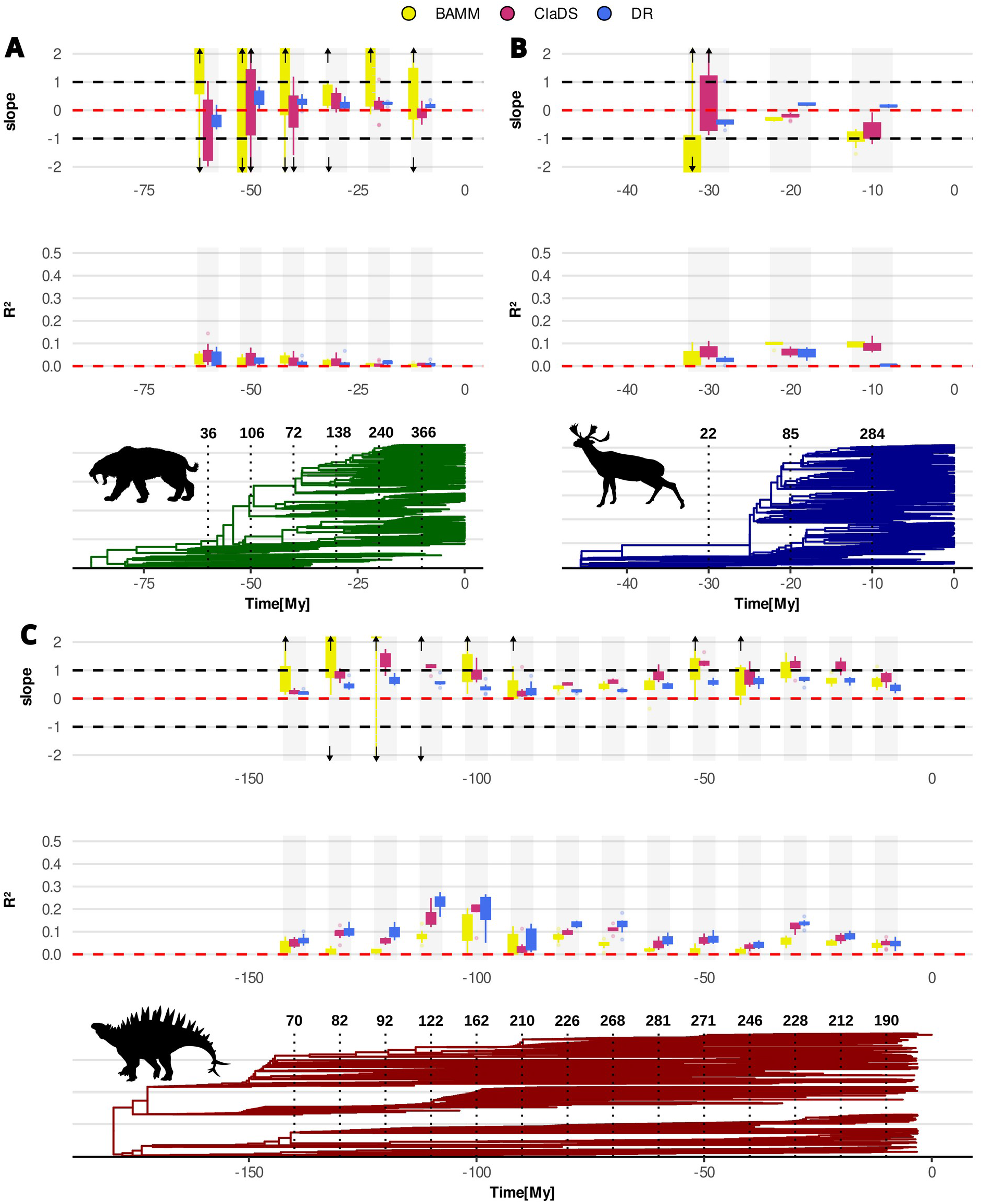
The relationship between tip estimates and subsequent rates of speciation (slopes) and the predictive capacities of tip estimates (R^2^), for BAMM (yellow), ClaDS (red), and DR (blue) at each slice in the fossil phylogenies of carnivorans (A), ruminants (B) and dinosaurs (C). Phylogenies were sliced every 10 million years and predictions were made for 5 million years from the slice. The numbers of tips at each slice are depicted in bold above the slice positions on the phylogenies. Note that 0 for dinosaurs refers to 66 My BP. The boxplots represent quartiles and 2*interquartile ranges across 10 samples from phylogenetic tree posteriors. The arrows indicate slope estimates lying outside the plotted range. The slope plots with full range of values are shown in Supplementary Figure 8.

### Predictive accuracy measures

We used the likelihood values of the BirDLinG models to calculate generalized R^2^, a measure commonly used in generalized linear models as an equivalent to Pearson’s coefficient of determination in normal linear models [23,24]. To calculate the generalized R^2^, it is necessary to compare the full model likelihood (*L_alt_*) with a null likelihood (*L_null_*) and a saturated likelihood (*L_sat_*). The null likelihood can be obtained using the same reasoning as the full model likelihood described above, but with *β_λ_* and *β_μ_* equal to 0 and only *α_λ_* and *α_μ_* optimised, i.e. with subsequent diversification rates *λ_subs_* and *μ_subs_*constant and independent of tip estimates *λ_tip_* and *μ_tip_*. The saturated likelihood is the maximum likelihood for each tip daughter species count *n* optimised individually, i.e. independently of the other tips in the dataset. This maximum is at

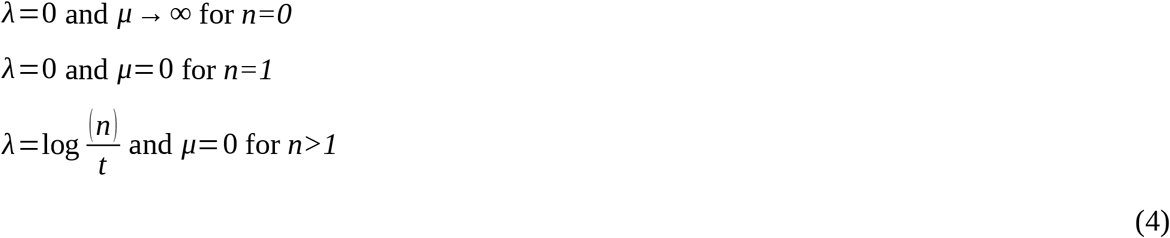

and equals the value of Equations 1 for fossil phylogenies or Equation 3 for extant phylogenies for respective *λ* and *μ* [21]. The total saturated likelihood is a product of the individual tip saturated likelihoods.

The generalized R^2^ using alternative, null and saturated likelihood is calculated as follows

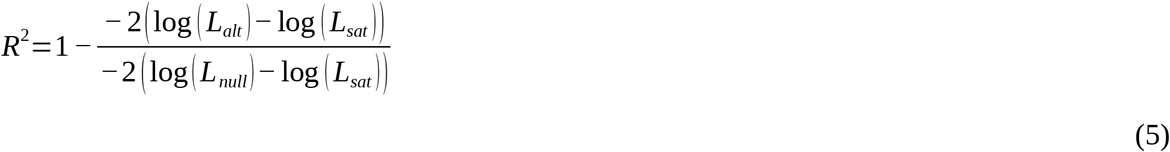

where the numerator and denominator of the fraction are deviances of alternative and null models, respectively.

Our formulation of generalized R^2^ allows us to tackle the information flow from tip rate predictions towards the number of subsequently surviving lineages in a statistically meaningful and comparable way; however, it cannot be interpreted as fully equivalent to the R^2^ of a normal linear model. Notably, even if the tip estimates of diversification were perfect predictors of subsequent rates, it would not lead to R^2^=1 [25], because the variance of the birth-death distribution depends non-trivially on both *λ* and *μ*. To ease the interpretation of R^2^ empirically obtained from extant phylogenies, we calculate the maximum achievable R^2^ on simulated sets with daughter species counts *n* drawn from a birth-death process with empirical *λ* and *μ* estimates obtained from ClaDS and BAMM (Supplementary Figure 4). A similar procedure was not possible for estimates based on DR metric due to the absence of extinction estimates from this method and also the observation that DR generally overestimates speciation rate (see below), which led to extremely large simulated trees in our pilot trials.

### Tip estimates of diversification rates

The tip rate estimation methods compared in this paper are BAMM [6], ClaDS [5] and the DR metric [4]. The BAMM model was fitted using the standard Metropolis coupled Markov chain Monte Carlo algorithm implemented in the BAMM procedure. The priors for each tree and time slice were generated with the function setBAMMpriors in the package BAMMtools [6]. Models were run with four chains at 100,000,000 iterations with 10% burn-in. Convergence was assessed with effective sample size (ESS >200) of the number of shifts. For the ClaDS model, we used its implementation in PANDA package in Julia (https://github.com/hmorlon/PANDA.jl), because it is considerably faster than the original R implementation. Models were run with three MCMC chains with 25% burn-in and otherwise default settings. Convergence was assessed after every 1000 iterations with the Gelman statistic [26], and the run was stopped once all the parameters had Gelman statistics<1.05. DR was calculated as the inverse of the Equal Splits measure [17] estimated using the R package picante [27]. The DR metric is proven to be an unbiased approximation of speciation rate in a pure birth process [4], so we used it as a tip estimate of speciation and considered tip estimates of extinction from DR to be 0. This effectively leads to cancelling parameter *β_μ_* from Equation 2. The BirDLinG models for DR were thus fitted with only constant extinction rate *α_μ_*, and no variation in extinction among the tips.

### Phylogenetic datasets

To test the predictive capacities of tip diversification metrics we used three phylogenetic datasets of tetrapod taxa including fossil tips (hereafter called fossil phylogenies). The phylogenies of ruminants (Ruminantia) and non-bird dinosaurs (Dinosauria) are taken from [13], and the phylogeny of carnivorans (Carnivora) comes from [28]. We limited our inference to these three fossil datasets because all the other available fossil phylogenies of tetrapods (e.g., in [13,29]) are too small for efficient tip rate inference (e.g., Testudinata, Equidae or Cetacea) and/or are taxonomic subsets of the dataset we use (e.g. Caninae within Carnivora). All trees are resolved at the species level and are significantly informed by fossil data where phylogenetic relations of fossil taxa are well established. The datasets contain both extant and extinct species except the dinosaurs which spans a period of 181.2 to 66 million years BP. We excluded the modern birds from the dinosaur phylogeny, including their stem lineage. We did this because there is no comprehensive fossilized phylogeny for birds after 66 My BP, and combining birds and dinosaurs would lead to a tree where a vast majority of the Mesozoic branchings are fossil-based and all the Cenozoic branchings are molecular-based. We however provide a separate analysis of molecular phylogeny of extant birds (see below). 10 trees were randomly sampled from a posterior sample of 100 trees for each dataset and used for downstream analyses. The fossil phylogenies were sliced every 10 million years and predictions were made 5 million years forward from every slice. This allowed us to identify possible differences in prediction ability due to the quality of more recent or ancient time scales while maintaining a consistent prediction window.

We also used extant molecular phylogenies of birds (Aves; [4]), amphibians (Amphibia; [30]), mammals (Mammalia; [31]) and squamate reptiles (Squamata; [32]). We chose these phylogenies because they are all relatively large (amphibians being the smallest, having molecular information on 4061 species; and birds being the largest, with molecular information on 6670 species) and they all have sampling fractions above 50%. Additionally we also analyzed separately the extant phylogenies of 6 large mammalian orders/suborders – ruminants (Ruminantia), carnivorans (Carnivora), primates (Primates), eulipotyphlans (Eulipotyphla), bats (Chiroptera) and rodents (Rodentia). These additional analyses allowed us to better interpret the trends observed in fossil ruminants and carnivorans, and also to explore the effect of phylogeny size on the quality of predictions. For all extant phylogenies, we randomly sampled 10 trees from the published posteriors to account for phylogenetic uncertainty. Extant phylogenies were sliced at 1, 5, and 10 million years before present and (because these trees do not include information on extinct branches, see above) predictions were made to the present.

## Results

### The relationship between tip and subsequent estimates of speciation and extinction

In order to understand the relationship between the tip estimates of diversification and the subsequent diversification rates (latently estimated based on observed species counts), we inspected the slopes and intercepts of the linear components of our BirDLinG model. For the speciation rates, the relationships between subsequent rates and tip estimates are generally positive, with some exceptions, for BAMM, ClaDS and also DR, for both fossil (Fig. 3 and Supplementary Figure 5 for aggregated histogram) and extant trees (Fig. 4 and Supplementary Figure 6 for aggregated histogram). This shows that all three methods can in principle serve as better than random predictors of speciation rates subsequent to the slice. Importantly, similar patterns are apparent using simpler Spearman correlations between tip rate estimates and subsequent numbers of species (Supplementary Figure 2 and 3), suggesting that this general result is independent of specific assumptions in BirDLinG. A more detailed look at the magnitude and variation among the individual time slices and taxa, however, reveals that the relationship between tip estimates and subsequent rates of speciation is fairly context dependent (Fig. 3 and 4).

**Figure 4:**
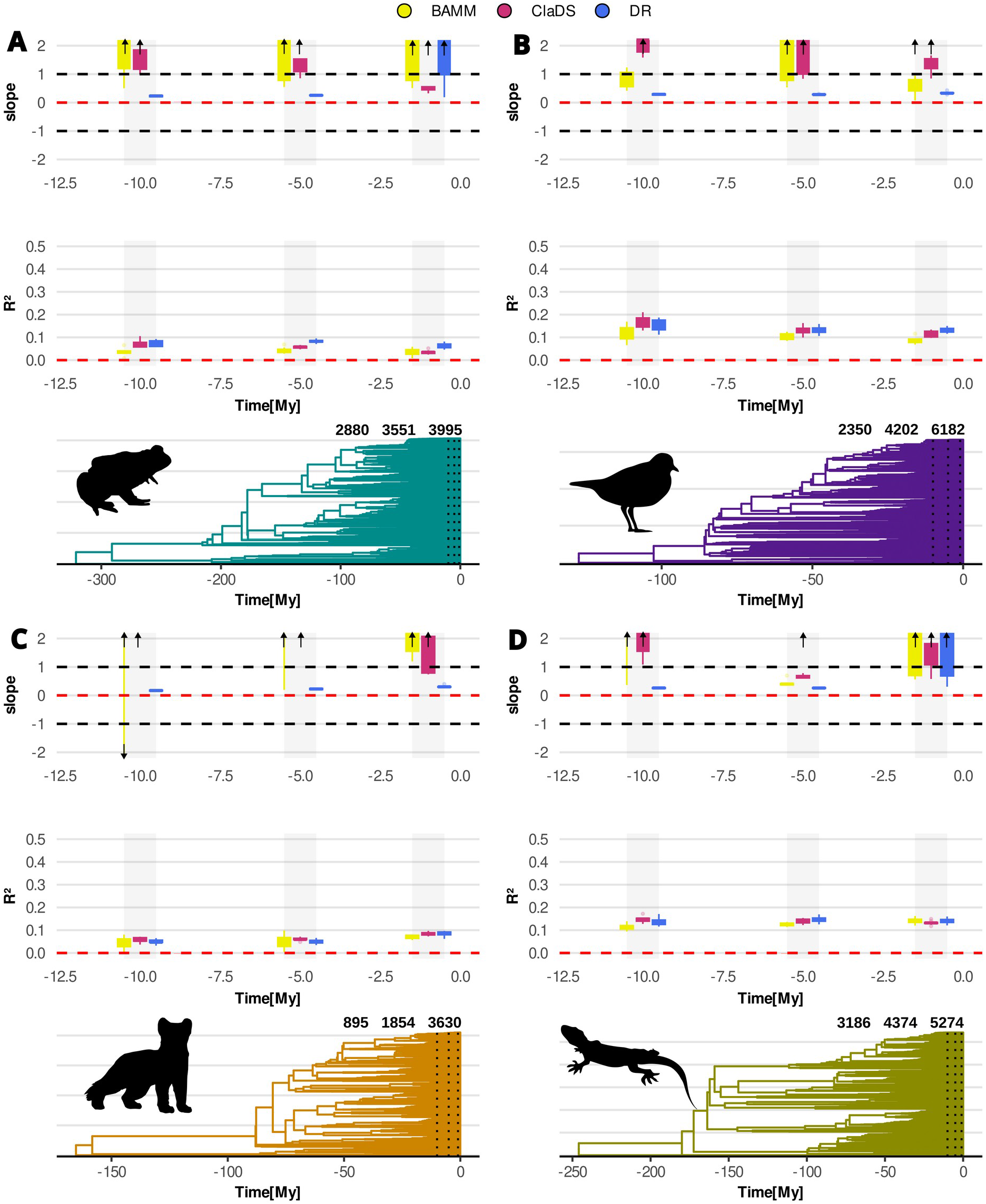
The relationship between tip estimates and subsequent rates of speciation (slopes) and the predictive capacities of tip estimates (R^2^), for BAMM (yellow), ClaDS (red), and DR (blue) at each slice in the extant phylogenies of amphibians (A), birds (B), mammals (C) and squamate reptiles (D). Phylogenies were sliced at 1, 5, and 10 million years before present and predictions were calculated to the present (i.e., the prediction windows are corresponding to 1, 5, and 10 million years). The numbers of tips at each slice are represented by bold triplets above the phylogenies. The boxplots represent medians, quartiles and 2*interquartile ranges across 10 samples from phylogenetic tree posteriors. The arrows indicate slope estimates lying outside the plotted range. The slope plots with full range of values are shown in Supplementary Figure 9.

Using fossil phylogenies (Fig. 3), the speciation slopes are generally positive for dinosaurs. For ClaDS and BAMM, they are fairly symmetrically scattered around the value of 1. This, together with speciation intercepts scattered around 0 (Supplementary Figure 7), shows that both methods retrieve correctly scaled estimates of speciation rates at the tips, athough with some spread and deviations for individual slices. The DR estimates for dinosaurs are also consistently positive, but are lower than 1. This shows that DR tends to predict higher variation of speciation rates than actually present in the subsequent diversification of dinosaurs. For carnivorans and ruminants, the speciation slope estimates using BAMM and ClaDS are often overlapping 0, showing that neither method works well for these phylogenies (Fig. 3). The most extreme examples are the slices of ruminants at 10 and 20 Ma BP, where the slopes are even consistently negative, i.e. the tips with highest speciation rate predicted by BAMM and ClaDS have the lowest subsequent speciation rates. In contrast to the two parametric methods, DR performs better on carnivoran and ruminant phylogenies, showing generally positive slopes, except for the 60 Ma BP slice for carnivorans and the 30 Ma BP slice for ruminants.

Across the extant datasets (Fig. 4), the speciation slopes are positive for all four used phylogenies, with exception of one tree sample of mammals and BAMM at 10 Ma BP. The slope estimates for BAMM and ClaDS tend to be scattered around 1, with a few outliers of very large values for some of the phylogenies and slices, especially for BAMM (Supplementary Figure 8 and 9). These outliers arose in cases where BAMM inferred no regime shifts, so that the subsequent speciation rate variation simply reflects the extremely conservative smooth component of the model. The slope estimates for DR are consistently larger than 0 but smaller than 1 (Fig. 4), and also the intercepts of DR are consistently positive for all four extant phylogenies except the 1 My BP slice in squamates (Supplementary Figure 10). The combintion of slopes between 0 and 1 and intercepts above 0 shows that the DR systematically underestimates the subsequent speciation rates by a constant but at the same time overestimates the variation of speciation rates across tips within each slice (see Supplementary Figure 11 for conceptual depiction).

The tip estimates of extinction, in contrast to speciation, do not appear to be meaningfully associated with subsequent extinction rates, and the slope estimates are distributed symmetrically around 0. This is as expected for extant phylogenies where the daughter species counts without zeros carry minimal information about subsequent extinctions (see Methods for details). However, this lack of association is true also for all the examined fossil phylogenies (Supplementary Figure 12), suggesting that neither BAMM nor ClaDS can correctly estimate extinction rate using phylogenetic trees. This is not surprising for ClaDS, where the extinction rates are simply the speciation estimates multiplied by a global extinction fraction parameter - this implementation is clearly meant only to control for presence of extinction in the system, not for estimating extinction rates. On the other hand, the tip extinction rates in BAMM are implemented using an independent process of rate evolution similar to one governing speciation, and could in theory serve for forecasting, although it is discouraged by the authors of the algorithm [15].

In order to interpret the lack of association between tip estimates and subsequent speciation in fossil phylogenies of ruminants and carnivorans (Fig. 3), we also analyzed extant-only trees of these two taxa and complemented them with four other mammalian orders of different sizes (Fig. 5). Our results suggest that DR tends to be positively associated with subsequent speciation in the majority of phylogenies and slices, with exception of the 10 Ma slices of ruminants and primates that contain around 30 and 50 tips, respectively, and also the 1 Ma slice for primates. In contrast, BAMM and ClaDS display a positive association only for 1 My slices of bats and rodents that contain more than 700 and 1400 species, respectively. The presented slices of fossil phylogenies of ruminants and carnivorans contain between 20 and 400 tips, suggesting that the lack of signal in these groups could be at least partially attributed to the size of these datasets.

**Figure 5:**
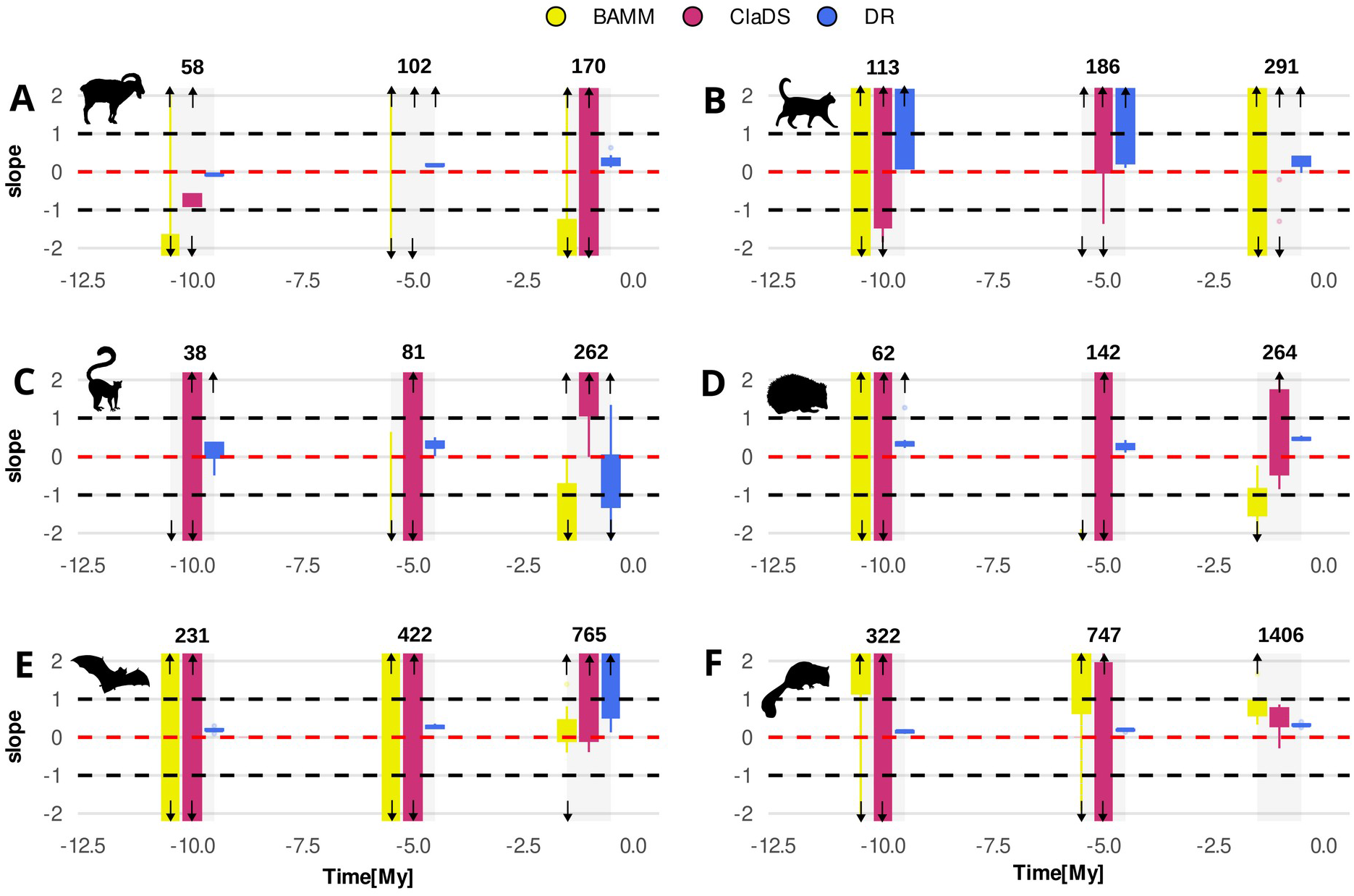
The relationship between tip estimates and subsequent rates of speciation (slopes) and the predictive capacities of tip estimates (R^2^), for BAMM (yellow), ClaDS (red), and DR (blue) at each slice in the extant phylogenies of ruminants (A), carnivorans (B), primates (C), eulipotyphlans (D), bats (E) and rodents (F). Like the four large extant phylogenies, the trees were sliced at 1, 5, and 10 million years before present and predictions were calculated to the present. The numbers of of tips at each slice are represented in bold above the plots. The boxplots represent medians, quartiles and 2*interquartile ranges across 10 samples from phylogenetic tree posteriors.

### Predictive capacity of tip rate estimates

We calculated generalized R^2^ based on our BirDLinG models as a measure of the tip rate estimates capacity to predict subsequent diversification. It is important to point out that the R^2^ values for fossil carnivorans and ruminants are generally not interpretable, because the speciation slopes are not positive, and the R^2^ thus reflects a spurious association. For the phylogenies with positive slopes, the R^2^ was typically between 0 and 0.1 for dinosaurs (Fig. 3 and Supplementary Figure 13 for aggregated histogram), amphibians and mammals, and between 0.1 and 0.2 for birds and squamates (Fig. 4 and Supplementary Figure 14 for aggregated histogram). There was no clear pattern of R^2^ with time, but ClaDS has generally slightly higher R^2^ than BAMM, and DR often outperforms both parametric methods.

The R^2^ in BirDLinG does not reach the value of 1 even under ideal circumstances, because mean and variance in the birth-death distribution depend non-trivially on both λ and μ. To ease the interpretation of empirical R^2^ in extant phylogenies, we calculated maximum achievable R^2^ for individual slices and methods based on a series of birth-death simulations using the empirical tip estimates for speciation and extinction. The maximum achievable R^2^ for ClaDS and BAMM were typically less than 0.5 (Supplementary Figure 4). The ratio between empirical and achievable R^2^ from simulations is typically around 0.5 for amphibians, birds and squamates, but lower for mammals. This suggests that the tip rate estimates can explain up to 50% of variation that is possible to explain accounting for the stochasticity inherent in the diversification process.

## Discussion

We have explored here the capacity of recently developed approaches for estimating tip rates of speciation and extinction (BAMM, ClaDS, and DR) to predict future diversification and diversity patterns. On average, all three methods produce tip estimates that are positively associated with subsequent speciation rates. This confirms that these methods are better predictors of future diversification than random guessing, a result that was previously demonstrated for DR on a more limited dataset of fossil-only phylogenies [13]. On the other hand, our results also suggest that the amount of variability in future diversity patterns that can be explained by tip rate estimates is fundamentally limited. The R^2^ values of empirical phylogenies were lower than 0.2, and the simulations suggest that much of the variation in subsequent diversity patterns is not deterministically predictable due to the large amount of stochasticity in the speciation process and overall unidentifiability of the extinction rates.

More worrying, although all the tip rate methods work satisfactorily on average, we found that certain combinations of methods and phylogenies produce estimates that are dissociated from subsequent dynamics, or even misleading. For instance in the phylogenies of carnivorans and ruminants, the BAMM and ClaDS tip estimates show no relationship with subsequent speciation dynamics, and in two slices of ruminants they are even negatively associated with subsequent speciation. There are two data-related artifacts that could explain this local underperformance: First, both carnivoran and ruminant phylogeny are a mixture of fossil- and molecular-reconstructed branches, leading to specific shapes of these trees. This explanation seems unlikely, because the lack of association between tip rates and subsequent speciation appears also in extant molecular-only phylogenies of carnivorans and ruminants (Fig. 5A and B). Second, BAMM and ClaDS fail on carnivorans and ruminants simply because these phylogenies are smaller than the rest of the datasets. Parametric methods like BAMM and ClaDS require relatively large phylogenies to converge [5,15]. While the slices of carnivorans and ruminants are between 20 and 400 species, which is commonly considered an adequate size for BAMM or ClaDS, they are still among the smaller phylogenies that we use in our study. The results for other mammalian orders of different sizes (Fig. 5) indeed suggest that the tip rate methods tend to underperform on smaller phylogenies. In particular, while DR estimates show a positive relationship with subsequent speciation, except for trees with very few tips, BAMM and ClaDS are only predictive for slices with circa 700 tips or more (1 My BP slices of rodents and bats).

In addition to these data artifacts, it is also possible that carnivoran and ruminant phylogeny truly exhibit patterns of speciation rate variation that cannot be captured under the assumptions of BAMM, ClaDS and DR. These methods expect that the speciation rates are locally positively correlated along the tree, but it is possible that carnivorans, ruminants and some of the other tested mammalian orders exhibit a lack of phylogenetic inertia or even negative correlation of rates across the trees. The latter could be caused by dynamics where past radiation causes lack of speciation in the future or decline of the clade. Such dynamics seem to be common in macroevolutionary systems [29] and can be caused by saturation of niche space [33,34], range size evolution [10] or various other processes [35]. Importantly, if the explanation through the absence of speciation heritability is correct, it very likely interacts with phylogeny sizes discussed in the previous paragraph – similarly as in linear regression, the quality of statistical signal in BirDLinG results from a combination of effect size and number of observations.

In the other phylogenetic datasets – dinosaurs, amphibians, birds, mammals and squamate reptiles – BAMM, ClaDS, and DR all give tip estimates of speciation that are generally positively associated with subsequent speciation rates. This suggests that an important component of speciation dynamics in these groups resembles the locally autocorrelated processes. Previous simulation results have shown that all three methods can recover speciation rates shifting according to a continuous Brownian motion, discrete rate-shift model, and several other locally autocorrelated mechanisms [5,15]. Interestingly, ClaDS and DR mildly outperform BAMM in prediction capacity in dinosaurs and the extant phylogenies. This might suggest that the modes of evolution with continuous or multiple small rate shifts as assumed by these methods are more represented in the empirical phylogenies than rarer large rate shifts as modelled by BAMM, but it might similarly reflect known problems with BAMM model implementation [36] or other issues. Another major difference among the methods is that DR shows a positive relationship with subsequent speciation more robustly than both ClaDS and BAMM. However, at the same time, DR also improperly scales with subsequent speciation, systematically underestimating it by a constant and at the same time overestimating the variation of speciation rates among the tips. This scaling pattern of DR is in accordance with previous findings via simulations [15], and is possibly related to the fact that both BAMM and ClaDS actively deal with extinction rates, whereas DR neglects extinction near the present [4]. Including extinction in models of diversification thus might be necessary for retrieving correctly scaled speciation estimates but comes at the expense of lower stability of the results.

The tip estimates of extinction rates seem generally unrelated to the expected values of subsequent extinction rates reconstructed from fossil phylogenies. This is not surprising - reconstructing extinction rates at the tips of molecular phylogenies is generally considered an extremely fraught, perhaps impossible, task [6,37,38]. Moreover, the response variable of our approach - the species counts - carries limited explicit information about subsequent extinction rates, as they are only indicated by the frequency of zeros in the slices of fossil phylogenies. On the other hand, the generally poor performance of the extinction estimation is the likely reason why we see no major qualitative differences between speciation estimation and predictive performances on fossil and extant phylogenies - the whole system carries so little information about extinction that formally conditioning the response variable on non-extinction does not cause any additional harm. Importantly, the comparable results of the analyses of the dinosaur and extant phylogenies suggest that our main conclusions are robust to the potential major sources of criticism of both data types, such as incomplete lineage sampling and general quality of the fossil data [19,20], as well as systematic omission of the lineages extinct after slicing in the extant phylogenies. This comparison suggests that, with an adequately conditioned model, the temporal slicing experiments can be to some extent performed on the molecular phylogenies without extinct species.

Our results have several interesting consequences for applications involving forecasting future diversification and diversity. We show that the tip rate estimates can, to a limited extent, inform us about future speciation potential (sensu [14]), even though they only inform us about background evolutionary rates, uninfluenced by human impacts (sensu [39]). On the practical side, we demonstrate that different methods of tip estimation have different properties and are potentially useful for different applications. In particular, DR is relatively robustly positively associated with subsequent speciation rates, but systematically overestimates their variation across the tips and underestimates their absolute value. DR is thus well tailored for tasks where preserving the order is more important than the the exact rate values, such as giving species conservation priorities based on future potential [13,14]. On the other hand, ClaDS and BAMM estimates are more unstable but centered around subsequent speciation rates. Estimates from these methods might be useful for comparing the current speciation rates based on phylogenetic trees with independent sources of evidence, such as the ones based on paleontological record [12,40], microevolutionary dynamics [41] or biogeographic processes [28,42].

An important problem with practical applications of tip rate metrics is that although they work on average better than chance, their statistical performance is widely variable based on individual phylogenies. In conservation applications, we may face a task where we have an extant phylogeny and want to predict future cladogenetic scenarios of individual species. Because we do not know the future we are predicting, we would have no direct way to know whether we work with a poorly predictable phylogeny, such as ruminants, allowing us to predict 1% of future diversification variability, or a phylogeny such as dinosaurs or birds allowing us to predict 10%. Luckily, our results show that the predictability is at least similar among neighbouring slices of one phylogeny. In this scenario, we thus recommend using BirDLinG on a few slices of the past of an extant phylogeny to gain a rough idea of its predictability in the future.

Predicting the future is always a complicated task, and predicting the future of a long term and extremely stochastic process, such as species diversification, is extremely challenging. In this paper, we map the limits of existing tip rate methods, associated with specific methodological assumptions, to predict future diversities and diversification dynamics. We show that the macroevolutionary dynamics can in principle be predicted, despite the modest statistical performance of the exising methods. The statistical performance can likely be improved by new macroevolutionary models and methods better incorporating the mechanisms of rate heritability and by critical benchmarking of these new models by their predictive capacity. In a broader perspective, we believe that the discussion about the predictive capacity of hypotheses and mechanisms is potentially an extremely stimulating future direction for macroevolutionary research. After all, predictions, both quantitative (e.g. species A will give rise to n daughter species) and qualitative (e.g., species are more likely to speciate if they are members of recently diversifying clade), are one of the most practical deliverables that macroevolution can provide to society and other scientific fields.

## Acknowledgements

We thank A. Toszogyová, I. Šímová, D. Pelletier, S. Lambert, S. Faurby, J. Cantalapiedra, D. Schluter and L. M’Gonigle for helpful comments and insights at different stages of the project. The research was funded by the Czech Science Foundation project GAČR 24-12851O to JS. AOM and SPO acknowledge NSERC Canada.

## Author contributions

GIR and JS conceived the study. JS designed the BirDLinG with assistance from SPO, AOM and DS, and GIR performed the evolutionary analyses. GIR and JS drafted the first version of the manuscript, and DS, AOM and SPO further edited it. All authors contributed to the final version of the manuscript and GIR and JS contributed equally.

## Code and data availability

The R package BirDLinG, pipelines used to analyze the phylogenies and primary phylogenetic trees are currently available in an anonymised FigShare repository https://figshare.com/s/f6e0ab1438fa07ff2ca1?file=70768068. In case of acceptance, BirDLinG will be linked to https://github.com/smyckaj/BirDLinG and the pipelines and data stored permanently.

