## Supplementary materials for "Phylogenies can be used to predict future diversification but with limited success"

[add author list in accepted version]

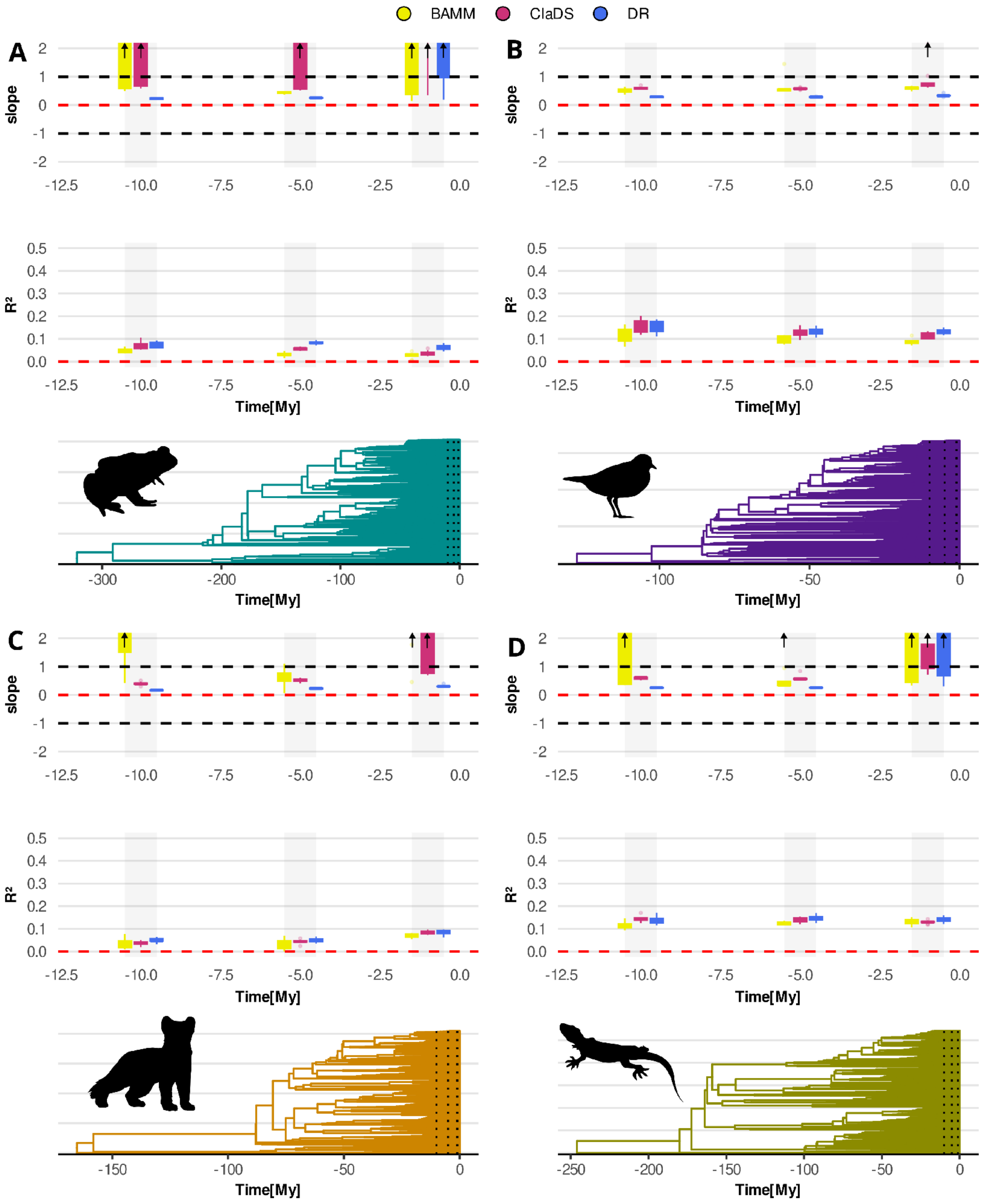


**Supplementary Figure 1:** Analysis of extant phylogenies with extinction slopes forced to 0. The relationship between tip estimates and subsequent rates of speciation are represented by the slope of the linear component of BirDLinG and the predictive capacities of different tip estimates are represented by $R^2$. The values are given for for BAMM (yellow), ClaDS (red), and DR (blue) at each slice in the extant phylogenies of amphibians (A), birds (B), mammals (C) and squamate reptiles (D). Phylogenies were sliced at 1, 5, and 10 million years before present and predictions were calculated to the present (i.e., the prediction windows are corresponding to 1, 5, and 10 million years). The boxplots represent medians, quartiles and 2*interquartile ranges across 10 samples from phylogenetic tree posteriors. The arrows indicate slope estimates lying outside the plotted range.

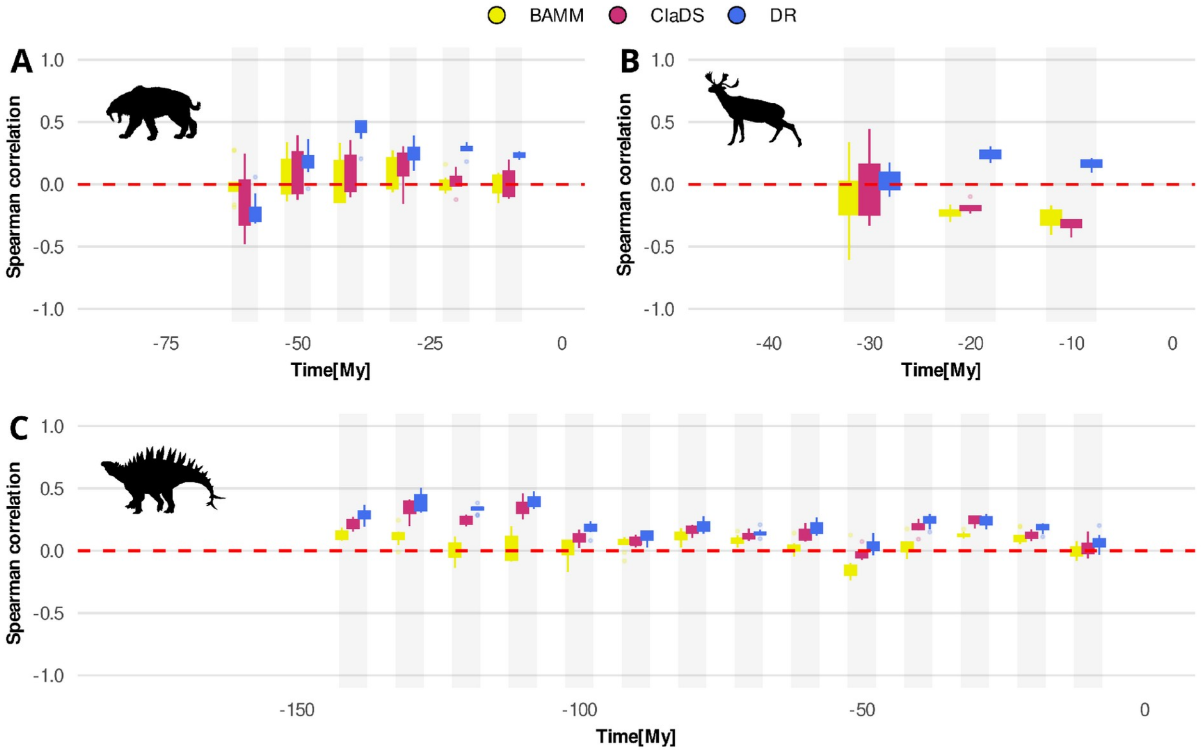


**Supplementary Figure 2:** Spearman correlation between tip speciation rates and subsequent species counts on slices of fossil phylogenies of carnivorans (A), ruminants (B) and dinosaurs (C). Phylogenies were sliced every 10 million years and predictions were made for 5 million years from the slice. The boxplots represent medians, quartiles and 2*interquartile ranges across 10 samples from phylogenetic tree posteriors.

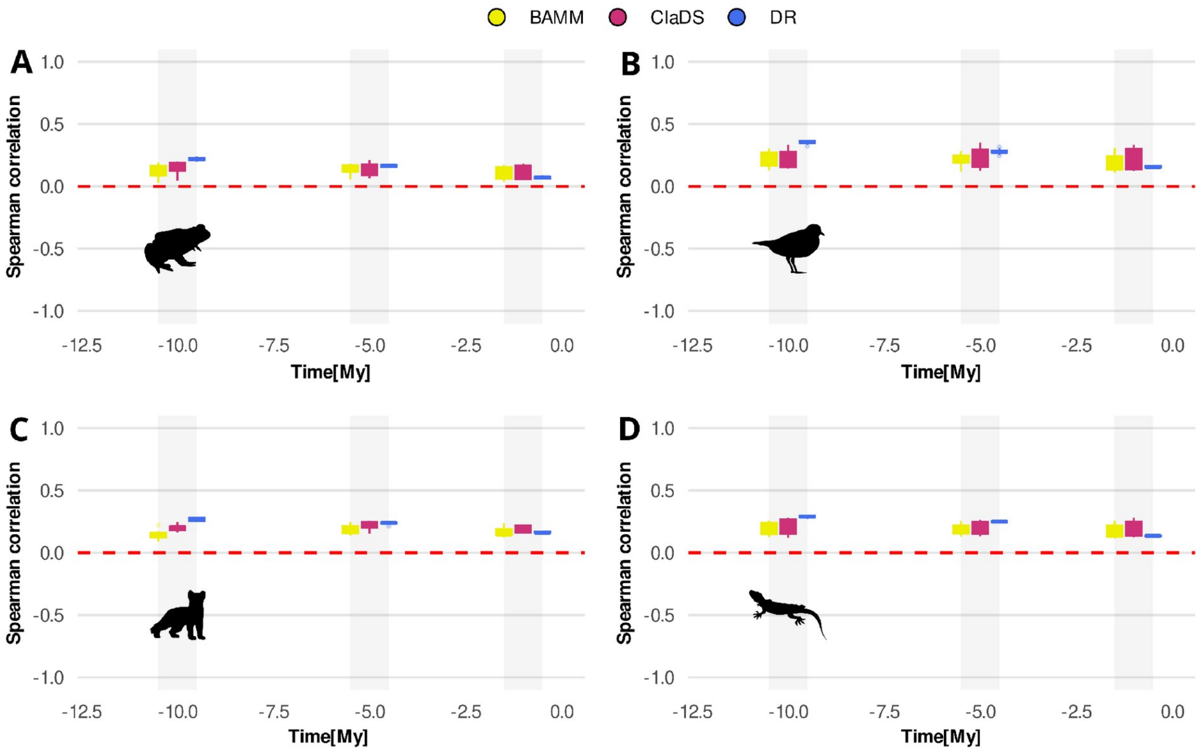


**Supplementary Figure 3:** Spearman correlation between tip speciation rates and subsequent species counts on slices of extant phylogenies of amphibians (A), birds (B) mammals (C) and squamate reptiles (D). Phylogenies were sliced at 1, 5, and 10 million years and predictions were calculated to the present. The boxplots represent medians, quartiles and 2*interquartile ranges across 10 samples from phylogenetic tree posteriors.

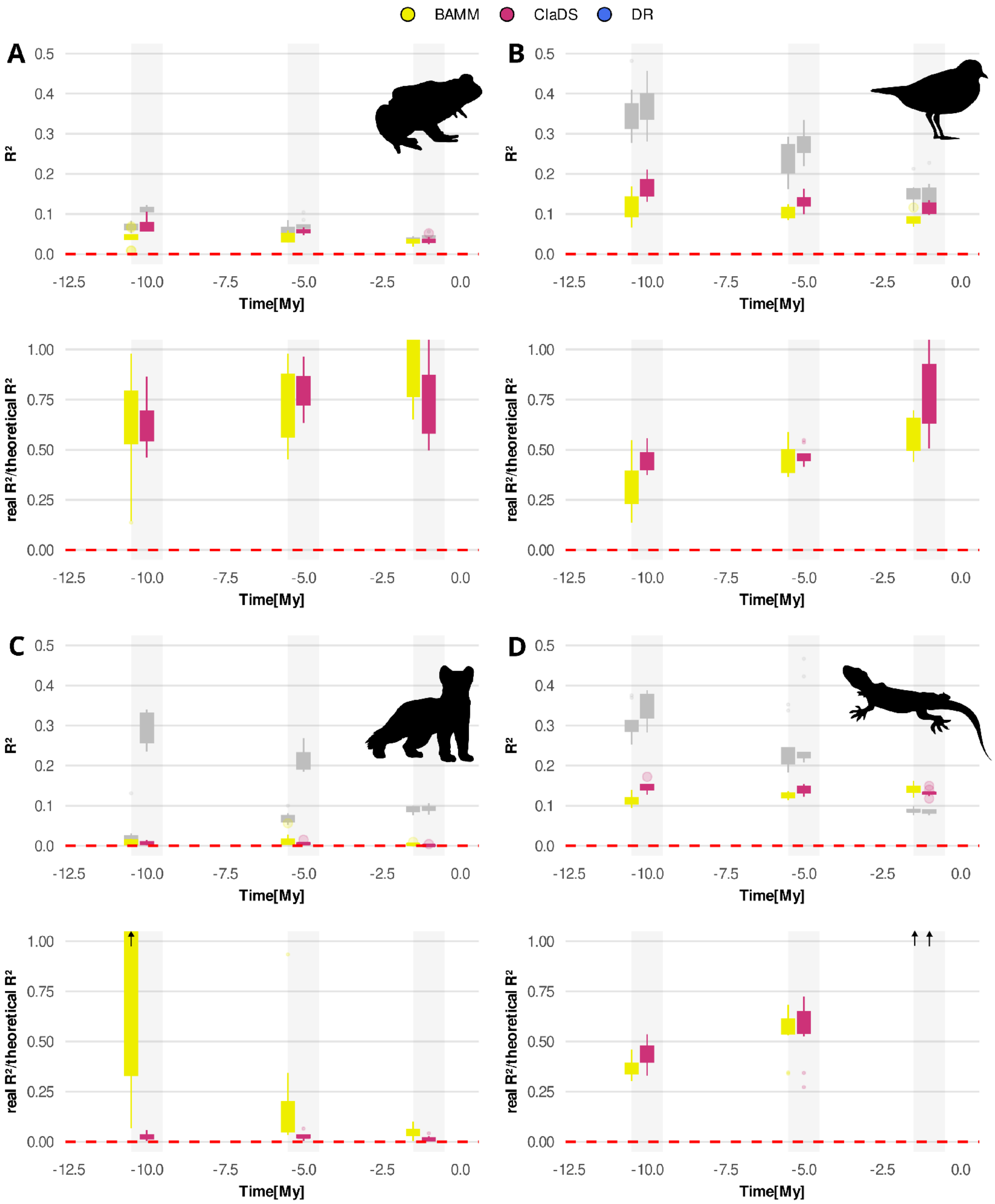


**Supplementary Figure 4:** Theoretical $R^2$ values based on daughter species counts simulated using tip rate estimates of BAMM and ClaDS for extant amphibians (A), birds (B), mammals (C) and squamate reptiles (D). The upper panel compares the achieved theoretical $R^2$ values (grey) with the empirical one (yellow, red), the lower panel depicts a ratio of empirical to theoretical $R^2$, Due to the stochastic nature of the simulations, the theoretical $R^2$ can be occasionally be lower than empirical one, especially in cases where empirical daughter species counts reach only few discrete values. The boxplots represent medians, quartiles and 2*interquartile ranges across 10 samples from phylogenetic tree posteriors. The arrows indicate slope estimates lying outside the plotted range.

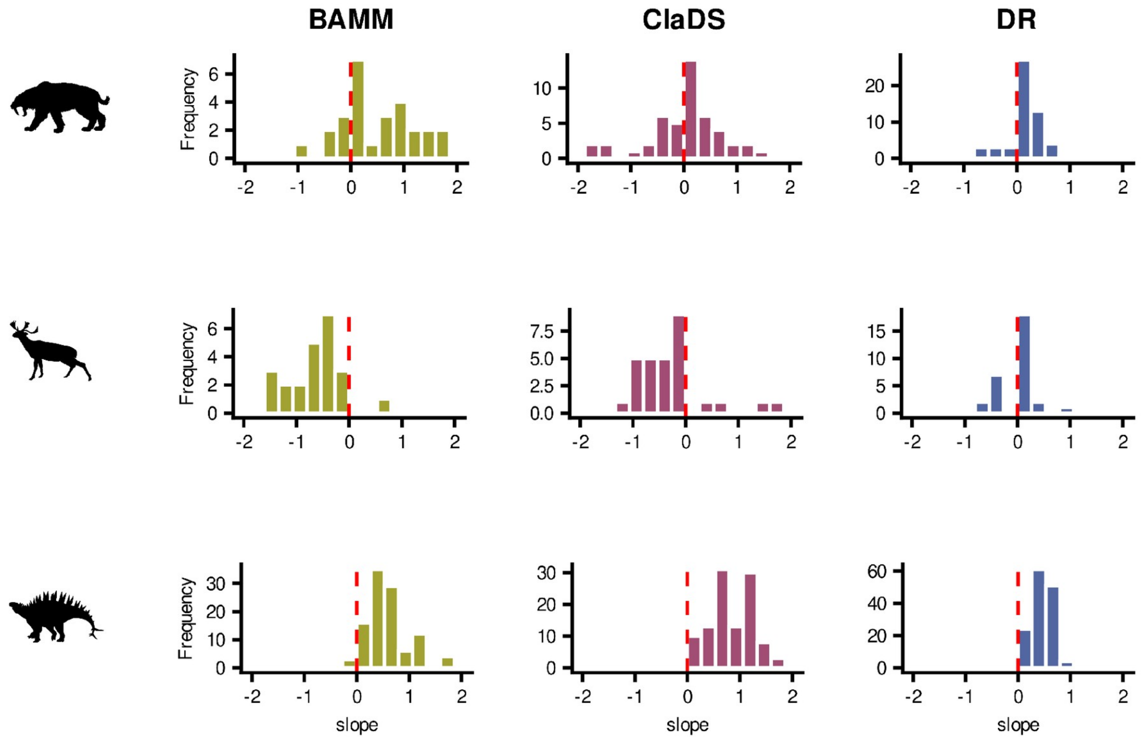


**Supplementary Figure 5:** Histograms of the regression slope relating subsequent rates of speciation and tip estimates for fossil phylogenies of ruminants, carnivorans. and dinosaurs collapsed across all timeslices. The range is adjusted to interval (-2, 2) for better readability.

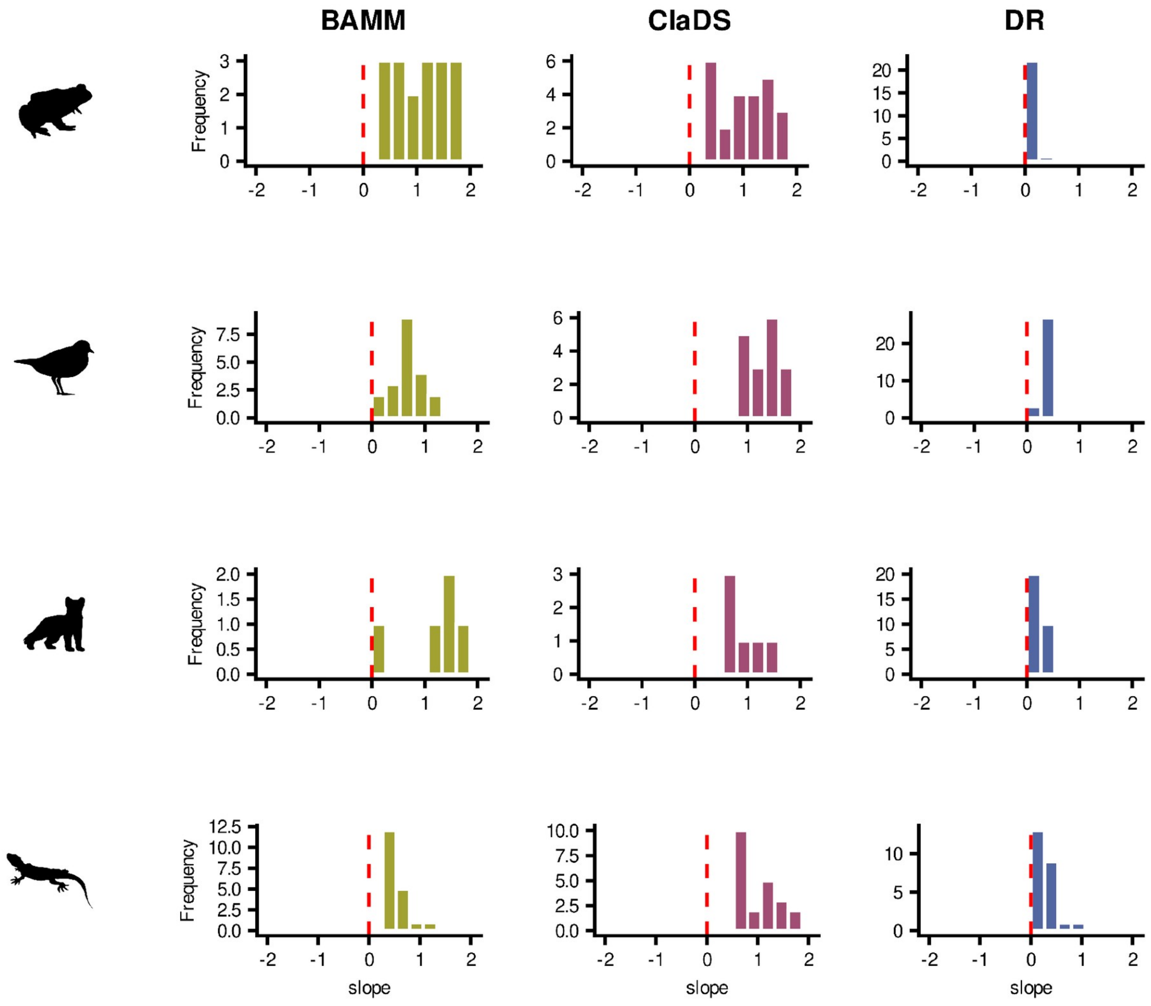


**Supplementary Figure 6:** Histograms of the regression slope relating subsequent rates of speciation and tip estimates for extant phylogenies of amphibians, birds, mammals and squamate reptiles collapsed across all timeslices. The range is adjusted to interval (-2, 2) for better readability.

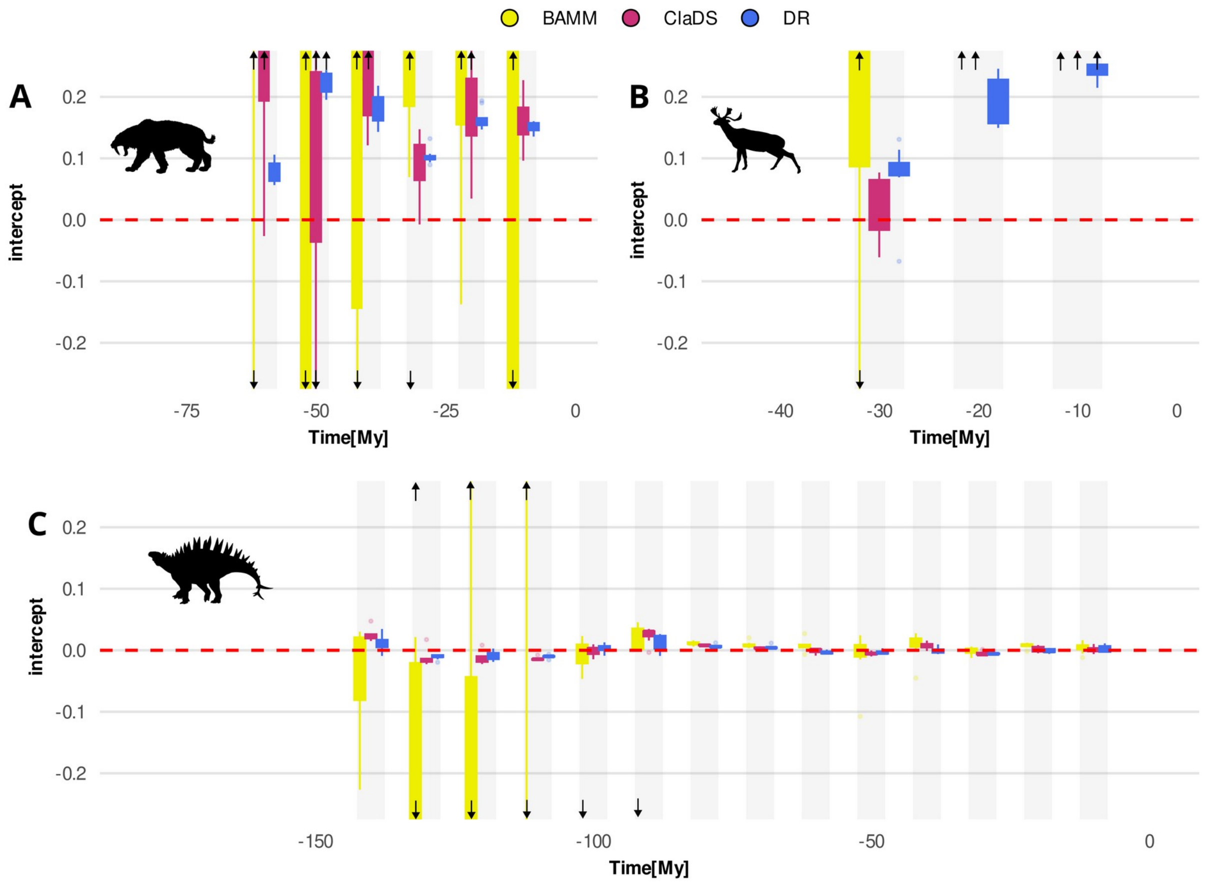


**Supplementary Figure 7:** The intercept of the linear relationships between tip estimates and subsequent rates of speciation for BAMM (yellow), ClaDS (red), and DR (blue) at each slice in the fossil phylogenies of carnivorans (A), ruminants (B) and dinosaurs (C). Phylogenies were sliced every 10 million years and predictions were made for 5 million years from the slice. Note that 0 for dinosaurs refers to 66 My BP. The boxplots represent quartiles and 2*interquartile ranges across 10 samples from phylogenetic tree posteriors. The arrows indicate intercept estimates lying outside the plotted range.

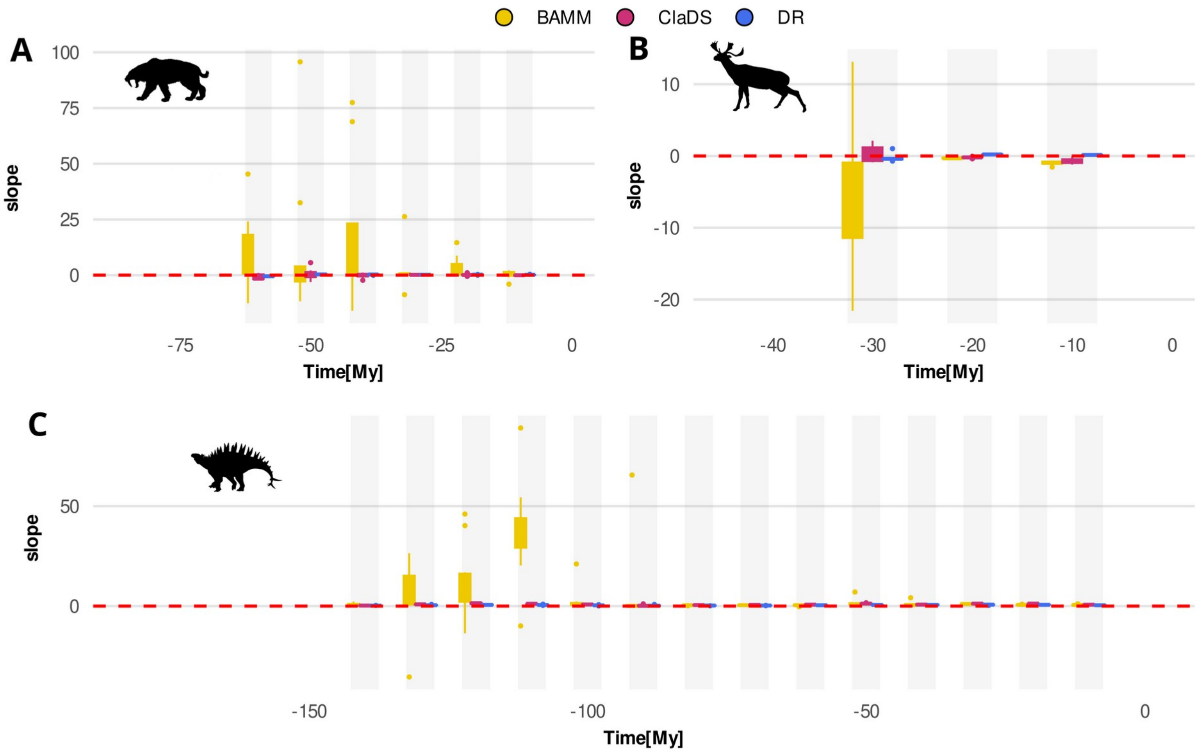


**Supplementary Figure 8:** The relationship between tip estimates and subsequent rates of speciation (slopes) for BAMM (yellow), ClaDS (red), and DR (blue) at each slice in the fossil phylogenies of carnivorans (A), ruminants (B) and dinosaurs (C). Phylogenies were sliced every 10 million years and predictions were made for 5 million years from the slice. Note that 0 for dinosaurs refers to 66 My BP. The boxplots represent quartiles and 2*interquartile ranges across 10 samples from phylogenetic tree posteriors. The arrows indicate slope estimates lying outside the plotted range. Unlike Fig. 3, the slope plots depict full range of values.

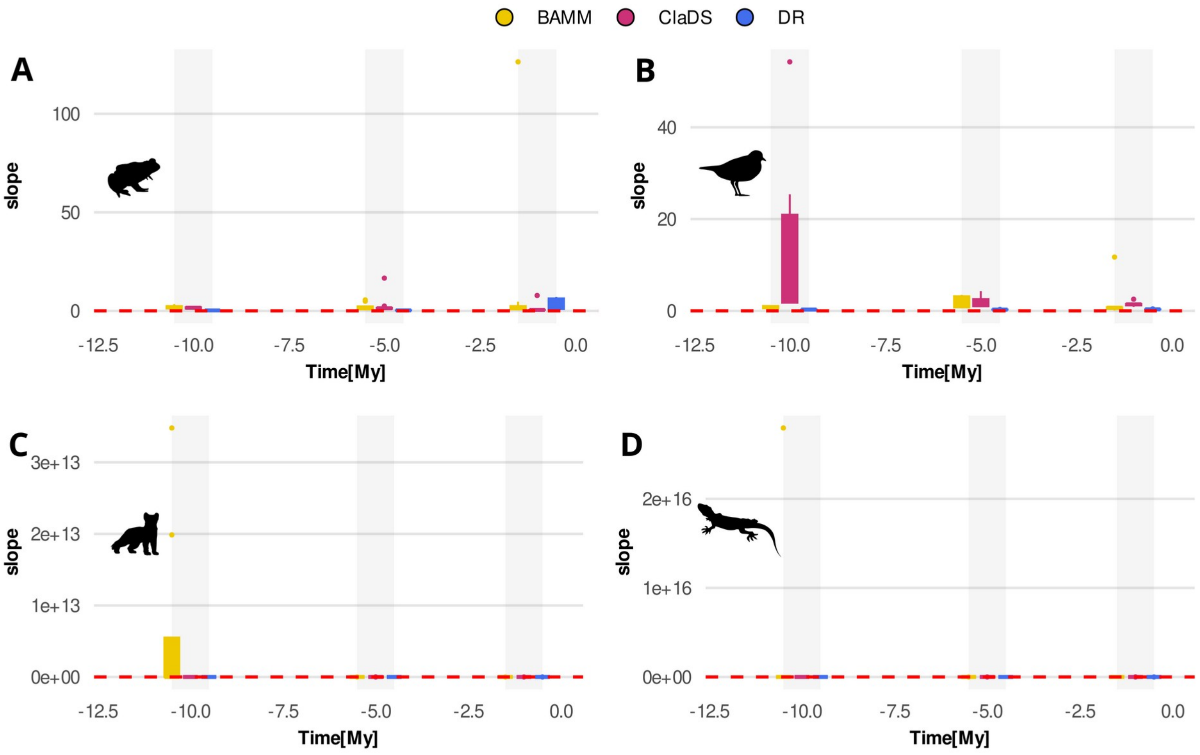


**Supplementary Figure 9:** The relationship between tip estimates and subsequent rates of speciation (slopes) for BAMM (yellow), ClaDS (red), and DR (blue) at each slice in the extant phylogenies of amphibians (A), birds (B), mammals (C) and squamate reptiles (D). Phylogenies were sliced at 1, 5, and 10 million years before present and predictions were calculated to the present (i.e., the prediction windows are corresponding to 1, 5, and 10 million years). The boxplots represent medians, quartiles and 2*interquartile ranges across 10 samples from phylogenetic tree posteriors.The arrows indicate slope estimates lying outside the plotted range. Unlike Fig. 4, the slope plots depict full range of values.

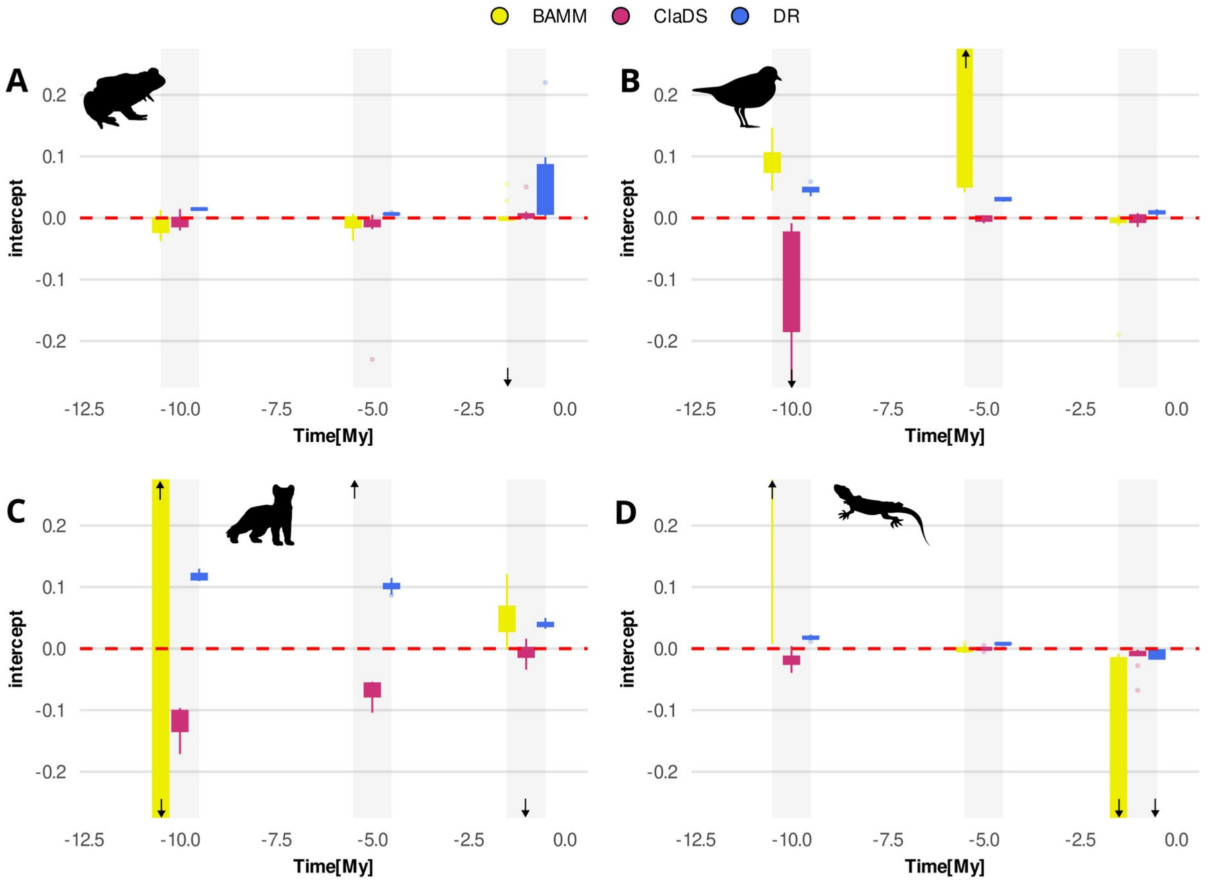


**Supplementary Figure 10:** The intercept of the linear relationships between tip estimates and subsequent rates of speciation for BAMM (yellow), ClaDS (red), and DR (blue) at each slice in the extant phylogenies of amphibians (A), birds (B), mammals (C) and squamate reptiles (D). Phylogenies were sliced at 1, 5, and 10 million years before present and predictions were calculated to the present (i.e., the prediction windows are corresponding to 1, 5, and 10 million years). The boxplots represent medians, quartiles and 2*interquartile ranges across 10 samples from phylogenetic tree posteriors. The arrows indicate intercept estimates lying outside the plotted range.

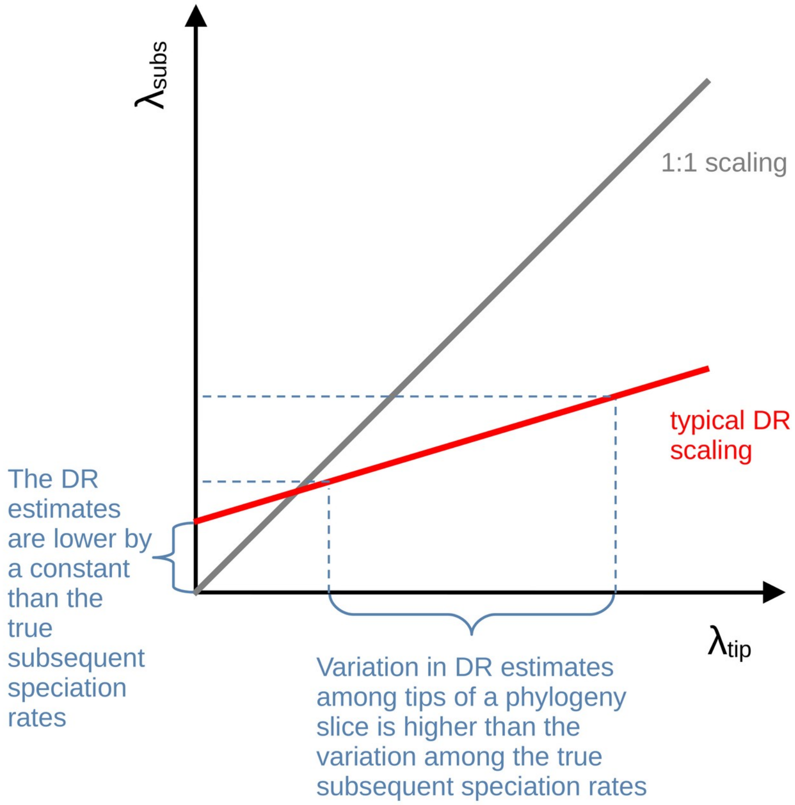


**Supplementary Figure 11:** The illustration of consequences of typical scaling of subsequent speciation rates ($\lambda_{subs}$) with the DR estimates ($\lambda_{tip}$), reflecting linear relationship with slopes typically falling between 0 and 1 and intercepts larger than 0.

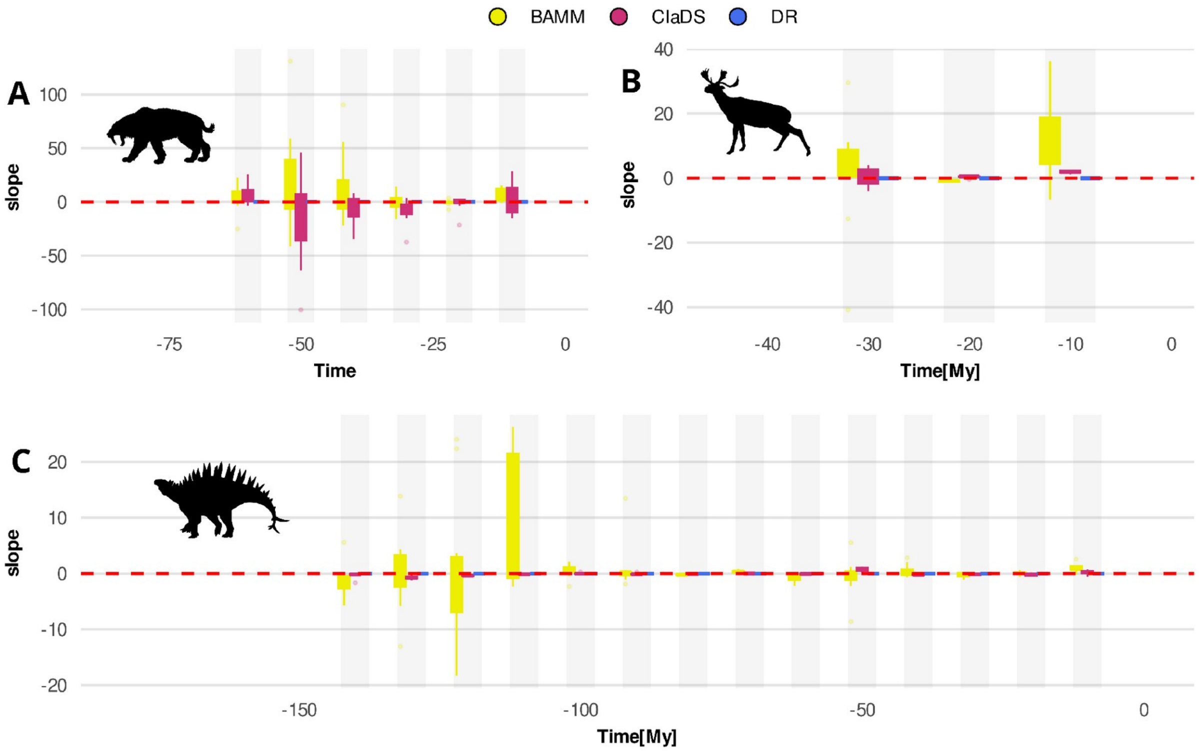


**Supplementary Figure 12**. The relationship between tip estimates and subsequent rates of extinction (slopes), for BAMM (yellow) and ClaDS (red). and DR (blue) at each slice in the fossil phylogenies of carnivorans (A), ruminants (B) and dinosaurs (C). The values for DR are equal to 0, because the metric does not provide any extinction estimate. Phylogenies were sliced every 10 million years and predictions were made for 5 million years from the slice. The numbers of tips at each slice are depicted in bold above the slice positions of the phylogenies. Note that 0 for dinosaurs refers to 66 My BP. The boxplots represent quartiles and 2*interquartile ranges across 10 samples from phylogenetic tree posteriors.

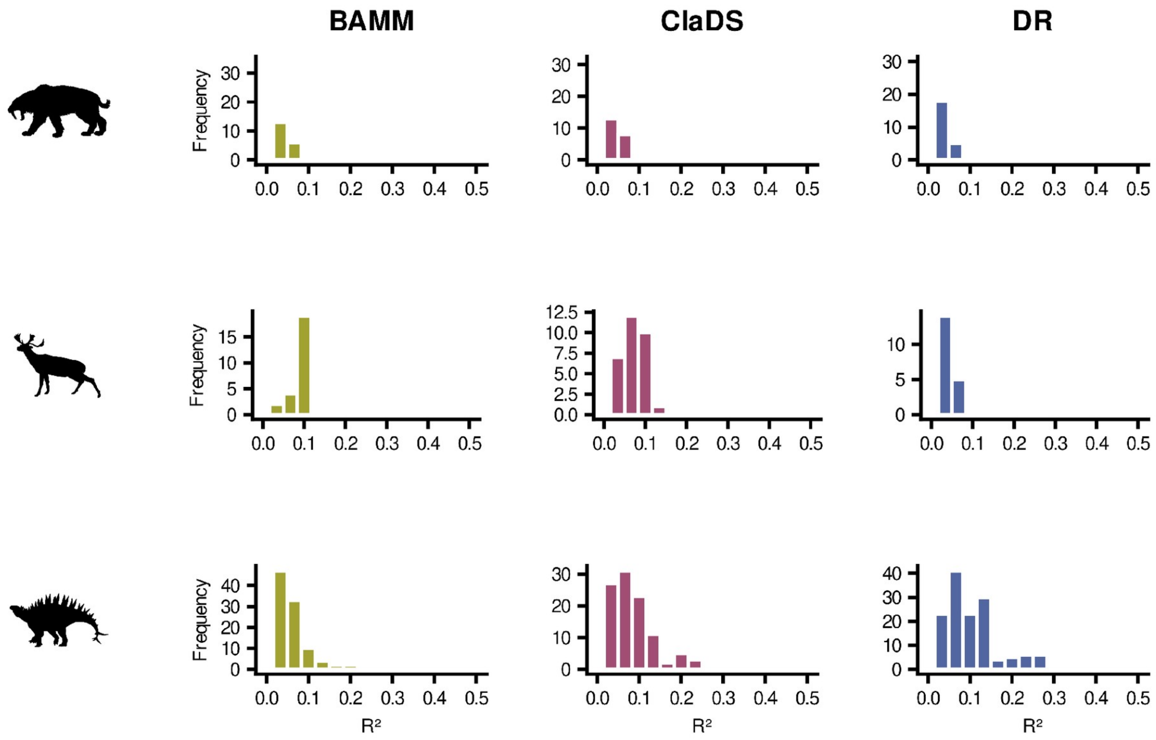


**Supplementary Figure 13:** The histograms generalized $R^2$ for fossil phylogenies of ruminants, carnivorans. and dinosaurs collapsed across all timeslices.

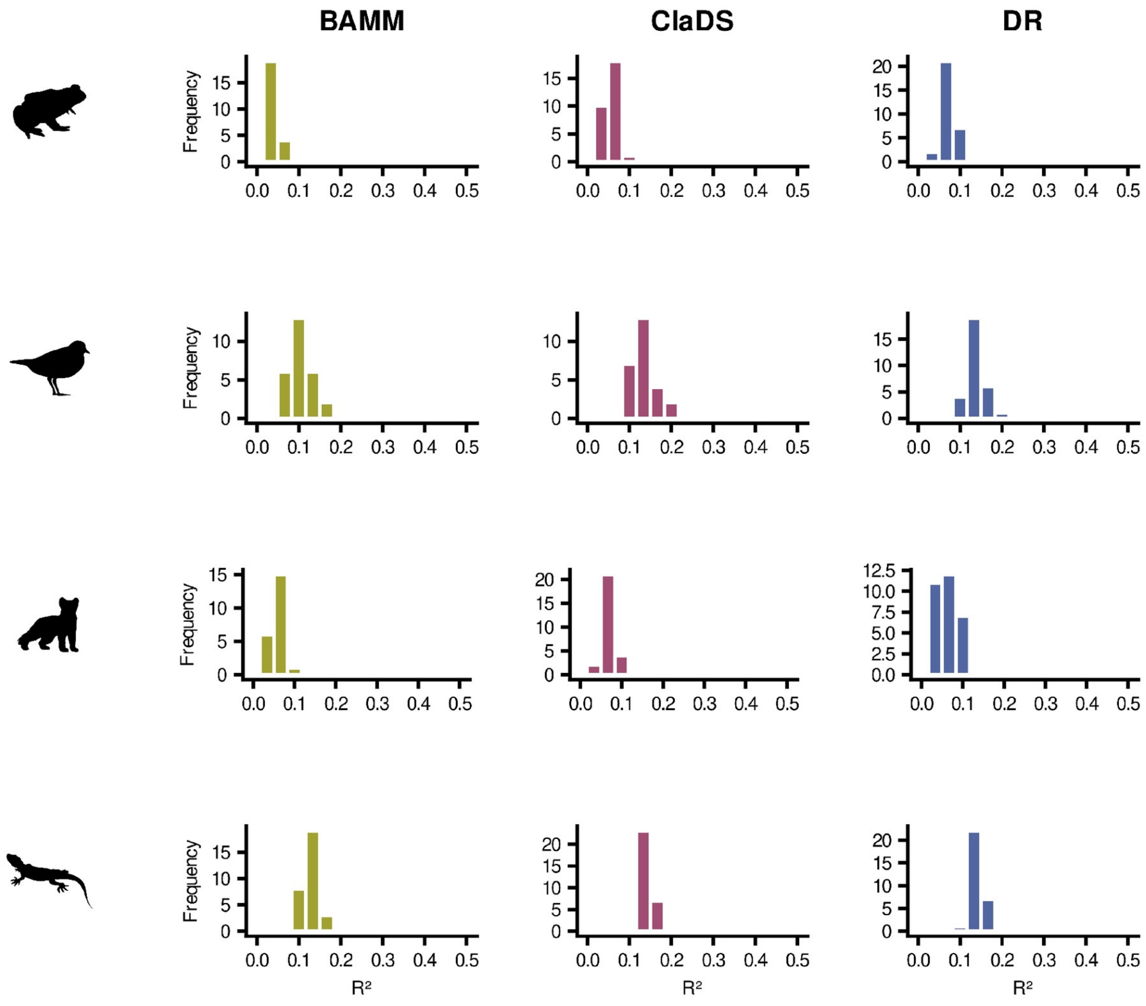


**Supplementary Figure 14:** The histograms generalized $R^2$ for extant phylogenies of amphibians, birds, mammals and squamate reptiles collapsed across all timeslices.
